# Allosteric regulation of switch-II controls K-Ras oncogenicity

**DOI:** 10.1101/2022.09.20.508702

**Authors:** Moon Hee Yang, Timothy H. Tran, Bethany Hunt, Rebecca Agnor, Christian W. Johnson, Timothy J. Waybright, Jonathan A. Nowak, Andrew G. Stephen, Dhirendra K. Simanshu, Kevin M. Haigis

**Author notes:** **Corresponding authors:** Kevin M. Haigis, Dana-Farber Cancer Institute, 450 Brookline Ave. LC-8312, Boston, MA, 02215., Dhirendra K. Simanshu, NCI RAS Initiative, Frederick National Laboratory for Cancer Research. 8560 Progress Drive, C1012, Frederick, MA 21701.

## Abstract

Ras proteins are GTPases that regulate a wide range of cellular processes. The activity of Ras is dependent on its nucleotide-binding status, which is modulated by guanine nucleotide exchange factors (GEFs) and GTPase-activating proteins (GAPs). Previously, we demonstrated that mutation of lysine 104 to glutamine (K104Q) attenuates the transforming capacity of oncogenic K-Ras by interrupting GEF induced nucleotide exchange. To assess the effect of this mutation *in vivo*, we used CRISPR/Cas9 to generate mouse models carrying the K104Q point mutation in wild-type and conditional K-Ras^LSL-G12D^ alleles. Consistent with our previous findings from *in vitro* studies, the oncogenic activity of K-Ras^G12D^ was significantly attenuated by mutation at K104 *in vivo*. These data demonstrate that lysine at position 104 is critical for the full oncogenic activity of mutant K-Ras and suggest that modification at K104, for example acetylation, may also regulate its activity. In addition, animals homozygous for K104Q were viable, fertile, and arose at Mendelian frequency, indicating that K104Q is not a complete loss of function mutation. Using biochemical and structural analysis, we found that the G12D and K104Q mutations cooperate to suppress GEF-mediated nucleotide exchange, explaining the preferential effect of K104Q on oncogenic K-Ras. Finally, we discovered an allosteric regulatory network consisting of K104 and residues including G75 on switch II (SWII) that is the key for regulating the stability of the α helix on SWII. In this allosteric network, K104-G75 interaction might be primary for keeping stabilization of SWII. Given the high frequency of KRAS mutations in human cancers, modulation of this network may provide a unique therapeutic approach.

## Introduction

K-Ras is a GTPase that regulate essential cellular processes by transducing extracellular signals emanating from cell surface receptors. Given the critical role of K-Ras in regulating cellular homeostasis, its functional activation – determined by its nucleotide binding state – is tightly controlled by two types of regulatory factors, guanine nucleotide exchange factors (GEFs) and GTPase activating proteins (GAPs) [1, 2]. GEFs activate K-Ras by facilitating the exchange of GDP for GTP, typically in response to external growth stimulatory signals. When GTP-bound, K-Ras can bind to effectors to transmit a proliferative signal or bind to GAP to promote GTP hydrolysis and functional inactivation [1, 2]. *KRAS* is among the most commonly mutated oncogenes, with missense substitutions at codon 12 (G12D is the most frequent) accounting for more than 70% of activating mutations [3]. Activating K-Ras mutations shift the steady-state equilibrium toward GTP-bound K-Ras by impairing intrinsic and GAP mediated hydrolysis [4, 5] and/or by increasing intrinsic and GEF mediated nucleotide exchange activity [6, 7].

Nucleotide binding controls K-Ras functional activity by influencing its conformational state, primarily at switch I (SWI; residues 30-38) and switch II (SWII: residues 60-76). Although both switch regions interact with GEF, GAP, and downstream effector proteins, SWI primarily facilitates GTP hydrolysis through GAP binding and SWII plays a significant role in binding to GEF to promote nucleotide exchange. Since most cancer associated K-Ras mutants sustain intrinsic GTPase activity [6], nucleotide exchange is still required for full activation. As such, modulating the equilibrium of the K-Ras-GTP/K-Ras-GDP cycle might be an effective therapeutic approach for targeting mutant K-Ras. Indeed, covalent inhibitors of K-Ras^G12C^ bind to the GDP-bound form and prevent nucleotide exchange [8].

In addition to regulation of its nucleotide binding state by GEFs and GAPs, K-Ras function is regulated by a variety of post-translational modifications (PTMs), including farnesylation, palmitoylation, phosphorylation, and ubiquitination, among others [9]. Previously, we identified lysine 104 (K104) as a site of K-Ras acetylation [10]. K104 is located in loop 7, between the α3 and β5 regions of K-Ras, which lies outside of the SWI and SWII domains [11, 12]. An acetylation-mimetic mutation of K104 (K104Q) decreased the activity of K-Ras^G12V^ in cellular transformation assays and molecular dynamics simulation suggested that K104 acetylation interferes with GEF mediated nucleotide exchange by destabilizing the SWII domain [10]. Here, we discovered allosteric network formed by interaction between K104 and three other residues including G75 on α2 helix in SWII and investigated a regulatory mechanism of perturbing this allosteric network by point mutation at K104 or G75 on oncogenic activity of K-Ras using multifaceted approaches.

## Results

### K104Q alters the oncogenic function of K-Ras^G12D^ in the intestinal epithelium

Previously, we demonstrated that a mutation that mimics acetylation at K104 attenuates the transforming capacity of oncogenic K-Ras *in vitro*, suggesting that it has a negative regulatory effect on K-Ras function [10]. Alternatively, K-Ras^K104Q^, like wild-type K-Ras, was found to rescue the growth deficit of cells lacking all Ras family members, suggesting that the mutation does not completely inhibit its function [13]. To address this discrepancy between these studies, we generated mouse embryonic stem cells in which the K104Q point mutation was inserted into the wild-type *Kras* gene and within a Cre-dependent activated allele using CRISPR/Cas9 technology (Fig. S1A) in mouse embryonic stem cells carrying a single copy of the *Kras*^*LSL-G12D*^ allele [14]. We subsequently used these cells to generate mouse models to study the effect of perturbing K104 in wild-type K-Ras and oncogenic K-Ras^G12D^ *in vivo*.

Expression of K-Ras^G12D^ in the intestinal epithelium promotes hyperplasia, but not neoplasia, by activating canonical MAPK signaling [15]. To investigate the effect of the K104Q mutation on the oncogenic activity of K-Ras, we crossed *Kras*^LSL-G12D;K104Q^ mice to *Fabp1-Cre* mice, which express Cre in the distal small intestinal and colonic epithelia [16]. At the histologic level, we observed that colons expressing K-Ras^G12D;K104Q^ exhibited a significant decrease in crypt height compared to those expressing K-Ras^G12D^ (Fig. 1A top panels and Fig. 1B). Consistent with this tissue-level phenotype, the number of proliferating cells on colonic crypts was significantly lower in the colonic epithelium expressing K-Ras^G12D;K104Q^ (Fig. 1A bottom panels and Fig. 1C). In addition to its effect on proliferation, activated K-Ras alters differentiation in the secretory cell lineage of the intestinal epithelium. This is most apparent in the small intestine, where epithelium expressing K-Ras^G12D^ lacks Paneth cells [15, 17]. Although clearly depleted, Paneth cells are present in small intestine expressing K-Ras^G12D;K104Q^ (Fig. 1D). Taken together, these data demonstrate that a secondary K104Q mutation can suppress the pro-proliferation and anti-differentiation phenotypes associated with a primary activating missense mutation in K-Ras.

**Figure 1.**
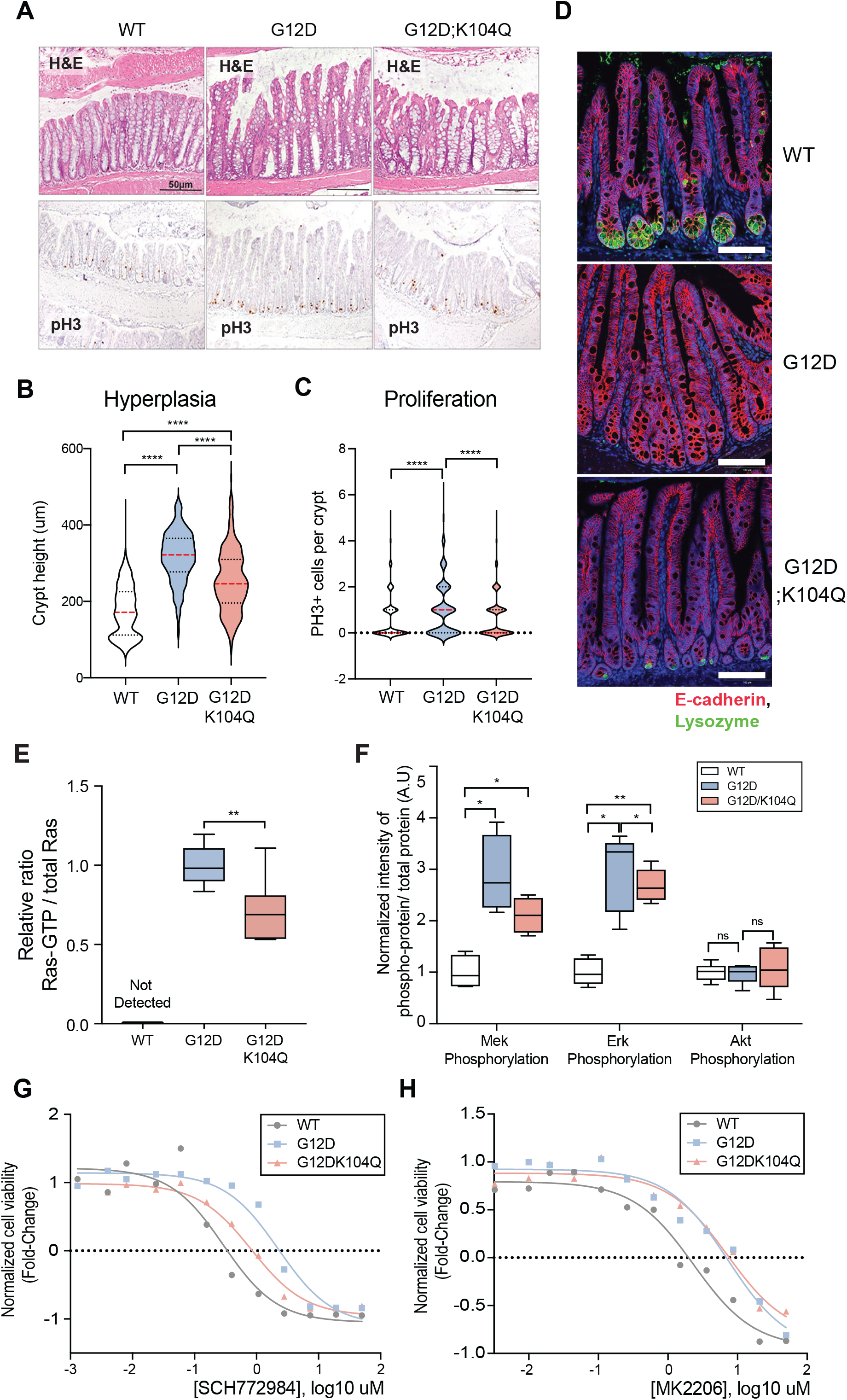
K104Q mutation negatively affects the K-Ras^G12D^ function in the intestinal epithelium through MAPK signaling pathway. (A) Histological analysis of the K-Ras wild-type and mutant colon. The top row shows representative H&E images of colonic epithelium from 8 - week-old *Fabp1-Cre* mice carrying the indicated *Kras* allele. The bottom row shows representative immunohistochemistry images for phospho-histone-H3 (pH3). Scale bars in all panels are 50m. (B) Quantification of crypt height in colonic epithelium. The K104Q mutation suppresses the hyperplasia phenotype associated with the expression of K-Ras^G12D^. All animals carried the *Fabp1-Cre* transgene and various alleles of *Kras*. Between 350-400 crypts from 4 different mice were counted for each genotype. Error bar shows SD. **** P < 0.0001, Mann-Whitney test. (C) Quantification of proliferation in colonic crypts. Secondary mutation, K104Q suppresses the hyperproliferative phenotype in colonic crypts expressing K-Ras^G12D^. Immunohistochemistry for pH3 was used to count the number of mitotic cells per crypt. Between 800-1000 colonic crypts from 4 different animals carrying the *Fabp1-Cre* transgene and *Kras* alleles were counted for each genotype. Error bar shows SD. **** P < 0.0001, Mann-Whitney test. (D) Staining small intestinal epithelium for Paneth cells. Representative immunofluorescence from *Fabp1-Cre* mice with indicated *Kras* genotypes for E-cadherin (magenta), a marker for epithelial cells, and lysozyme (green), a marker for Paneth cells. Scale bars indicate 100 µm. The K104Q mutation weakly suppresses the effect that K-Ras^G12D^ has on the secretory cell lineage differentiation. (E) Ras activation in colons expressing different K-Ras mutants. The levels of Ras-GTP were assessed using Raf-RBD Pull-down assay. K104Q mutant reduces K-Ras activity in colons expressing K-Ras^G12D^. N=9 for WT, G12D, and G12D;K104Q in 3 independent experiments. The error bar shows SD. ** P < 0.005 (F) Activation of canonical Ras signaling pathways in colons expressing different K-Ras mutants. Elevated MAPK signaling activation in colons expressing K-Ras^G12D^ is suppressed by a second site mutation, K104Q. Activation of MAPK signaling was measured by western blotting for Mek phosphorylation at Ser217/221 and Erk phosphorylation at Thr202/204. PI3K signaling activation was determined by western blotting for Akt phosphorylation at Ser473. N=6 for WT, G12D, and G12D;K104Q. Error bar shows SD. * P < 0.02; ** P < 0.005, Mann-Whitney test. (G-H) Response of colonic organoids carrying different *Kras* mutant alleles to inhibiting MAPK or PI3K signaling pathway by specific inhibitor treatment. SCH772984 and MK2206 were used for inhibiting Erk and Akt respectively. K-Ras^G12D^ organoids are resistant to inhibiting Erk and Akt relative to ones expressing wild-type K-Ras. K104Q mutant confers the sensitivity to Erk inhibitor treatment, but not to Akt inhibition in K-Ras^G12D^ organoids. Each dot in inhibitor response curve represents the average of quadruplicate and curves were fit with nonlinear regression.

Mutationally activated K-Ras promotes proliferation in the colonic epithelium through canonical MAPK signaling [15, 18, 19]. To determine how the K104Q mutation alters K-Ras^G12D^ signaling, we measured the level of GTP-bound K-Ras^G12D^ by quantifying its interaction with the Ras binding domain (RBD) of Raf-1 in protein lysates from the colons. The strong K-Ras activation seen upon expression of K-Ras^G12D^ in the intestinal epithelium was decreased by the K104Q mutation (Fig. 1E and Fig. S2A). Consequently, canonical MAPK signaling – measured by immunoblotting for phosphorylated Mek and Erk1/2 – is also decreased in colons expressing K-Ras^G12D;K104Q^ relative to those expressing K-Ras^G12D^. However, PI3K signaling, another canonical K-Ras effector, is not changed in colons expressing either K-Ras^G12D^ or K-Ras^G12D;K104Q^ (Fig. 1F and Fig. S2B). This decrease in MAPK signaling is clearly not related to reduced steady state expression of K-Ras^G12D;K104Q^, since it is expressed at the same level as K-Ras^G12D^ (Fig. S1C) supported by marginal impact of K104Q on protein thermostability (Fig. S5A).

Furthermore, we determined the effect of K104Q mutant on oncogenic K-Ras^G12D^ in colonic epithelium organoids system. K-Ras^G12D;K104Q^ organoids show attenuated proliferation compared to K-Ras^G12D^ organoids (Fig. S3A), demonstrating that this system recapitulates our *in vivo* observations. Consistently, the robust activation of MAPK signaling by K-Ras^G12D^ is decreased in K-Ras^G12D;K104Q^ organoids (Fig. S2B). Interestingly, while K-Ras^G12D^ organoids are resistant to ERK and AKT inhibition K104Q sensitizes organoids to ERK inhibition, but not to PI3K inhibition (Fig. 1G and 1H). These results are consistent with a model in which K104Q decreases the phenotypic output of K-Ras^G12D^ by affecting its steady-state level of GTP loading, which translates into reduced downstream signaling through MAPK.

### K104Q negatively impacts oncogenic K-Ras in colon cancers

Although mutationally activated K-Ras is sufficient to alter colonic epithelial homeostasis through its effects on proliferation and differentiation, it is not sufficient to promote tumorigenesis and therefore requires cooperation with loss-of-function mutations in tumor suppressor genes [15]. To determine whether K-Ras^G12D^ is still sensitive to the suppressive activity of K104Q mutation in colon tumors, we crossed *Fabp1-Cre* mice carrying *Kras*^LSL-G12D;K104Q^ to those carrying a Cre-dependent conditional allele of the Adenomatous polyposis coli (*Apc*) tumor suppressor gene [20]. Strikingly, we observed that the number of the tumors was significantly reduced in animals with tumors expressing K-Ras^G12D;K104Q^ relative to those expressing K-Ras^G12D^ and, in fact, the number of colon tumors that developed in the background of K-Ras^G12D;K104Q^ expression was not significantly different from when Apc-mutant tumors expressed only in wild-type K-Ras (Fig. 2A). Consistent with this result, the overall survival of animals bearing K-Ras^G12D;K104Q^ colon tumors was significantly extended compared to those bearing K-Ras^G12D^ colon tumors (Fig. 2B). Previous studies suggest that Wnt signaling cooperates with activation of MAPK signaling by K-Ras mutation [21, 22]. We measured the level of Wnt signaling by assessing the nuclear β-catenin on tumors of varying *Kras* genotypes. Tumors expressing K-Ras^G12D^ has higher levels of nuclear β-catenin, and it is decreased in ones expressing K-Ras^G12D;K104Q^ (Fig. 2C and 2D). However, despite significant differences in survival and tumor number, tumor morphologies for different *Kras* genotypes were all characterized as adenomas with low-grade dysplasia (Data not shown).

**Figure 2.**
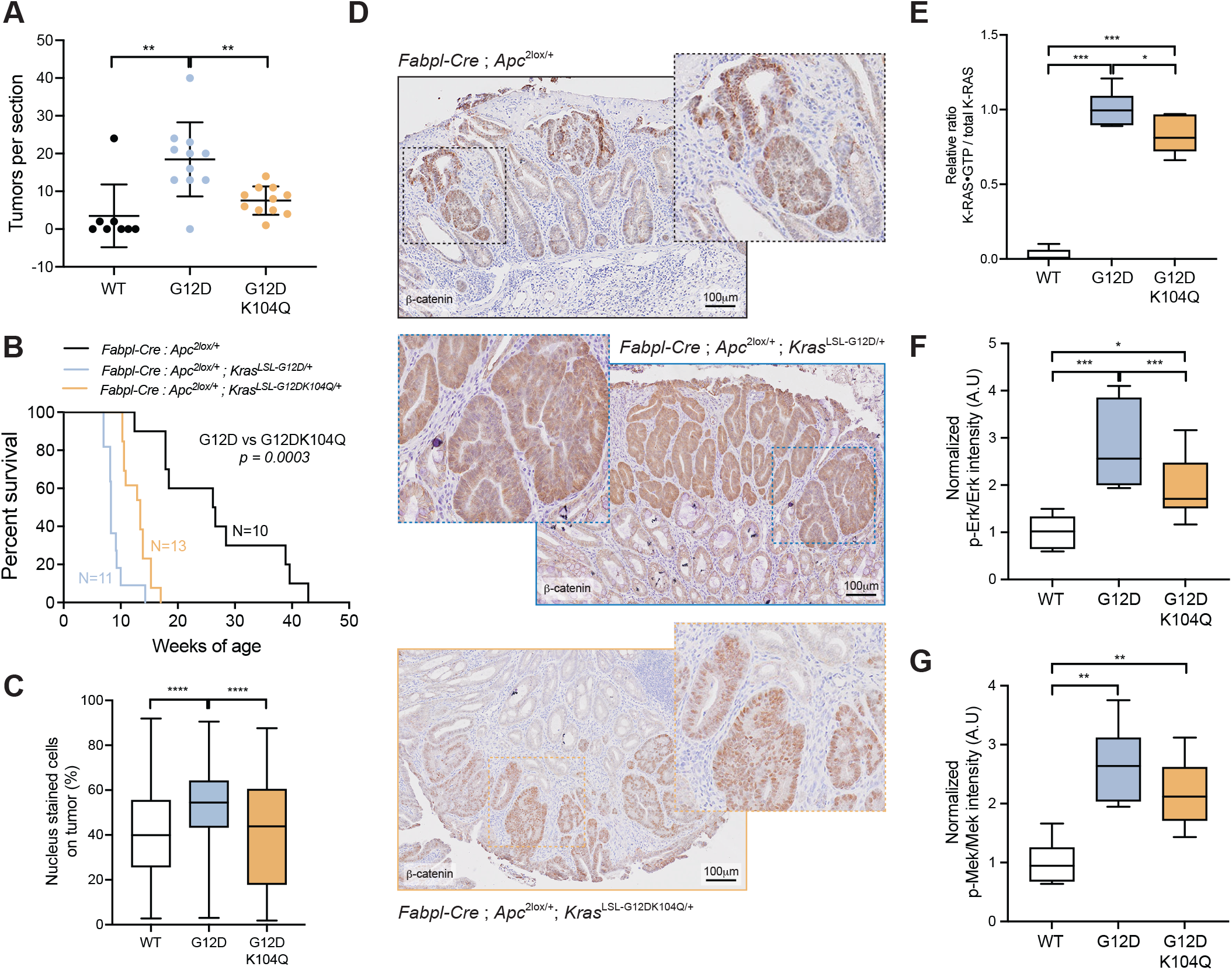
The effect of K104Q mutation on the colon tumor model. (A) Scatter plot demonstrating the number of dysplastic lesions in animals carrying the indicated genotypes. The number of dysplastic lesions in colon tumor model expressing K-Ras^G12D^ is significantly reduced by secondary mutation, K104Q. N=8 for *Fabpl-Cre; Apc*^*2lox/+*^, N=12 for *Fabpl-Cre; Apc*^*2lox/+*^; *K-Ras*^*LSL-G12D/+*^, and N=10 for *Fabpl-Cre; Apc*^*2lox/+*^; *K-Ras*^*LSL-G12D;K104Q/+*^. ** P < 0.002, Mann-Whitney test. (B) Survival curves for *Fabpl-Cre; Apc*^*2lox14/+*^ animals bearing colorectal tumors. Each animal carried the indicated *Kras* allele. P value was calculated using log-rank test. (C) Representative immunohistochemistry for -catenin on colon tumors expressing different K-Ras mutants. Scale bars in all panels indicate 100 µm. (D) Graph illustrating quantification of nuclear β-catenin on colon tumors expressing different K-Ras mutants. Oncogenic K-Ras enhances the nuclear localization of β-catenin in colon tumors, and it is weakly reduced by secondary mutation at K104. Between 80-240 dysplastic lesions from 4-5 different mice were counted for each genotype. Error bar shows SD. (E) Activation of K-Ras in colon tumors carrying different *Kras* alleles. K-Ras activation was assessed using Raf-RBD pull-down assay. The elevated Ras activity is slightly decreased by K104Q mutant in colon tumors expressing K-Ras^G12D^. N=12 for WT and G12D;K104Q and N=8 for G12D in 3 independent experiments. (F-G) Activation of MAPK, a canonical K-Ras signaling pathway, in colon tumors carrying different *Kras* mutant alleles. Graphs illustrated the quantification of western blotting analysis for Erk (F) and Mek [49] activation. MAPK signaling pathway is highly activated by expressing K-Ras^G12D^. K104Q mutant significantly decreases the activation of Mek and Erk in colon tumors carrying oncogenic mutant *Kras*. N=5 mice per genotype in 2 independent experiments. In all panels, error bar shows SD. * P < 0.02; ** P < 0.002; **** P < 0.0001, Mann-Whitney test.

As in our study of colons expressing K-Ras, but wild-type for Apc, we sought to understand how the secondary K104Q mutation affects the biochemical properties of K-Ras^G12D^. In tumors expressing K-Ras^G12D;K104Q^, the level of GTP-bound K-Ras was slightly, but significantly, decreased compared to those expressing K-Ras^G12D^ (Fig. 2E and Fig. S4A). As in our studies on the non-neoplastic colonic epithelium, Erk activation was significantly reduced (Fig. 2F and Fig. S3B), but Mek activation was decreased but not significantly (Fig. 2G and Fig. S3C) in tumors expressing K-Ras^G12D;K104Q^. Altogether, our studies demonstrate that perturbing lysine 104 of K-Ras – in this case by mutating it to glutamine – negatively impacts the oncogenic output of K-Ras^G12D^, likely through canonical MAPK signaling in both intestinal epithelium and colorectal tumors.

Additionally, we determined the effect of K104Q on wild-type K-Ras in our *in vivo* model. Because constitutional disruption of the *Kras* gene causes embryonic lethality secondary to failure of fetal liver and cardiovascular development at embryonic day 12.5 [23, 24], we intercrossed mice carrying the *Kras*^*K104Q/+*^ allele with those carrying *Kras*^*LSL-G12D;K104Q/+*^, which functions as a null allele in the absence of Cre to determine whether the K104Q mutation suppresses the developmental function of K-Ras. All genotypes of offspring from this cross were born at the expected Mendelian frequency (Table1). *Kras*^*K104Q/LSL-G12D;K104Q*^ mice were phenotypically indistinguishable from *Kras*^*+/+*^ mice. Moreover, livers from e12.5 embryos heterozygous or homozygous for *Kras*^*K104Q*^ did not show significant changes in the expression or activation of Ras compared to wild-type controls (Fig. S1B). Collectively, these observations demonstrate that a single allele of *Kras*^*K104Q*^ is sufficient to support normal embryogenesis indicating that it is not a loss-of-function mutant consistent with previous findings *in vitro* [13].

### K104Q modulates nucleotide exchange activity through Switch II conformation

Because K104Q mutant regulates the oncogenic activity of K-Ras by modulating the level of GTP-bound Ras directly *in vivo* (Fig. 1E & 2E), we determined the effect of K104Q on K-Ras GDP-GTP equilibrium cycle by measuring GTP hydrolysis and nucleotide exchange. We observed no effect of K104Q mutation on the NF1-mediated GTP hydrolysis in wild-type K-Ras and K-Ras^G12D^, suggesting that reduced oncogenicity of the K-Ras^G12D;K104Q^ is not due to restoration of GAP-mediated GTPase activity (Fig. 3A). Previously, Yin. G. *et al*. showed a 2-fold reduction in p120-RasGAP-mediated GTPase activity for the K-Ras^K104Q^ mutant [13]. This small difference in GAP activity could be due to the different RasGAPs (NF1 vs. p120-RasGAP) and their concentration (1 nM NF1 vs. 50 nM p120-RasGAP) used in the assay. Next, we assessed the effect of K104Q on nucleotide exchange and found that K104Q synergistically inhibits both intrinsic and SOS catalyzed nucleotide exchange (Fig. 3B). This observation suggests that G12D may rely on SOS for full activation *in vivo*, and a plausible rationale for why the presence of K104Q mutation in wild-type K-Ras has a milder effect on its activation.

**Figure 3.**
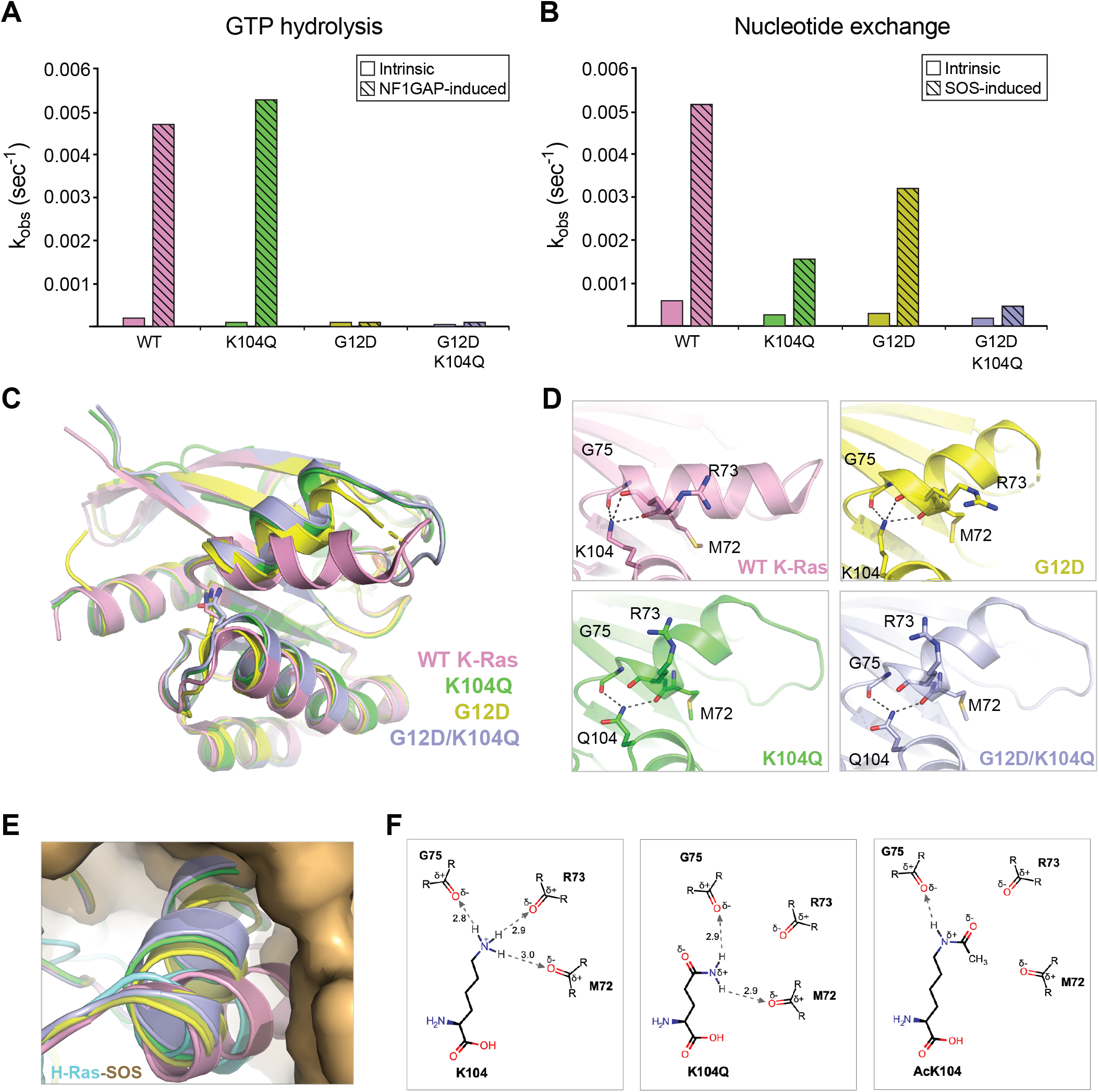
K104Q mutation in oncogenic K-Ras^G12D^ suppresses GEF-mediated nucleotide exchange. (A) Bar graph showing the rate of intrinsic and NF1-mediated GTP hydrolysis for WT K-Ras, K-RasK104Q, K-RasG12D, and K-RasG12D;K104Q. In this assay, phosphate ions released by GTP hydrolysis from GTP-loaded K-Ras protein were captured by a phosphate-binding protein containing a phosphate sensor present in the reaction buffer. The fluorescence signal from the phosphate sensor during the capture of phosphate ions was monitored over time in the absence (intrinsic) and presence of NF1-GAP. (B) Bar graph showing intrinsic and SOS-mediated nucleotide exchange rate for wild-type and mutant K-Ras (K-Ras^K104Q^, K-Ras^G12D^, and K-Ras^G12D;K104Q^). The rate of GDP release from mant-GDP-loaded K-Ras protein was determined by monitoring the decrease in mant-GDP fluorescence signal as a function of time in the absence (intrinsic) and presence of SOS1. (C) Overlay of crystal structures of GDP-bound K-Ras (pink, PDB:6MBT), K-Ras^K104Q^ (green), K-Ras^G12D^ (yellow, PDB:5US4), and K-Ras^G12D;K104Q^ (light blue). Residue K/Q104 and GDP are shown as sticks for all the structures. (D) Enlarged view of the interactions formed by residue K/Q104 with M72, R73, and G75 of a2 helix is shown for each of the four structures shown in panel C using the same color-coding scheme. Relevant side chains are shown as sticks with hydrogen bonds drawn as dashed lines. (E) Enlarged view of the H-Ras-SOS interface with structures of wild-type K-Ras, K-Ras^K104Q^, K-Ras^G12D^, and K-Ras^G12D;K104Q^ superposed on H-Ras. The α2 helix in Ras proteins is shown in cartoon representation, whereas SOS is shown in surface representation. Structural superposition shows the stearic clash of a2 helix present in the SII region of K-Ras^K104Q^, K-Ras^G12D^, and K-Ras^G12D;K104Q^ with SOS at the Ras-SOS interface. (F) Schematic representation of interactions formed by K104, K104Q, and acetylated K104 with M72, R73, and G75 residues presenting in the α2 helix. Most robust interactions are formed by K104 in wild-type K-Ras structure which involves hydrogen bonds, and partial electrostatic interactions with M72, R73, and G75 (left panel). Q104 residue in the K-Ras^K104Q^ structure retains hydrogen bonds and electrostatic interactions with M72 and G75 but unable to interact with R73 (middle panel). A structural model of acetylated K104 in K-Ras predicts that acetylated lysine would only be able to interact with G75 and not with M72 and R73 (right panel).

To understand the structural basis for the function of K104Q on GEF mediated nucleotide exchange of K-Ras, we crystalized and solved structures of K-Ras^K104Q^ and K-Ras^G12D;K104Q^ in the GDP-bound state at resolutions of 1.59 Å and 1.84 Å, respectively (Table S1). Superimposed, all four K-Ras structures share the same overall fold, but there is a significant shift in the helix of SWII region of K104Q, G12D and G12D;K104Q mutants (Fig. 3C and 3D). In the wild-type K-Ras structure, K104 helps stabilizing the α2 helix of SWII by interacting with the main-chain carbonyl oxygen atoms of M72, R73 and G75 (Fig. 3D, top left panel). The side chain atoms of K104 also exhibit the same set of interactions in the K-Ras^G12D^ structure (Fig. 3D, top right panel). In the structures of single mutant (K104Q) or double mutant (G12D:K104Q) of K-Ras, the amide group present in the side chain of Q104 is constrained to a planar geometry and because of that forms H-bonds with M72 and G75, but not R73. Loss of the K104-R73 interaction shifts the switch II helix relative to wild-type K-Ras (Fig. 3D, bottom panels). Interestingly, a similar but lesser shift in switch II is seen in the K-Ras^G12D^ structure, perhaps underlying the cooperation seen between G12D and K104Q mutations in our nucleotide exchange experiments (Fig. 3B and 3D).

Our biochemical data suggest that the interactions formed by K104 and residues M72, R73 and G75 in SWII have a significant role in GEF binding to RAS, and that reorientation of SWII is enough to destabilize this interaction. Based on the structural superposition of K-Ras structures described above with the previously solved structure of the H-Ras-SOS complex [25], a change in the orientation of the α2 helix of SWII in the structures of K-Ras^K104Q^, K-Ras^G12D^, and K-Ras^G12D;K04Q^ would clash with SOS at the Ras-SOS interface (Fig. 3E), providing a structural explanation as to why this mutation leads to decreased rates of nucleotide exchange. Furthermore, the shift in SWII induced K104Q mutation may selectively inhibit the K-Ras-GEF interaction. Evaluation of Raf and Pi3k binding to K-Ras using isothermal titration calorimetry (ITC) assay showed that K104Q had no effect on interaction of both downstream effectors with either wild-type K-Ras or K-Ras^G12D^ (Fig. S5B and S5C).

### Perturbation of the SWII allosteric network

We initially generated the K104Q mutation to mimic the constitutive acetylation of K-Ras, although computational simulation predicted that K104 acetylation would have a greater effect on K-Ras structure than K104Q mutation [10]. Considering our structural analysis, we re-addressed this prediction by examining how acetylated lysine would interact with M72, R73, and G75 residues, which together with K104 form an allosteric network controlling the orientation of the α2 helix in SWII. Modeling of acetylated lysine in K-Ras structures suggests that acetylated lysine forms a single hydrogen bond with the carbonyl oxygen of G75 (Fig. 3F). The partial negative charge on the oxygen atom present in the acetyl group would repel the main-chain carbonyl oxygen of R73. The methyl group of acetylated lysine points towards the carbonyl oxygen atom of M72, resulting in loss of interaction between acetylated lysine and M72. A comparison of lysine, glutamine, and acetylated lysine at the 104 positions in K-Ras suggests that acetylated lysine would destabilize the SWII helix to a greater extent than the K104Q mutation (Fig. 3F). In all cases G75 retains the interaction with both K104-Ac or K104Q, indicating that this interaction is required to be conserved.

To gain insight into the function of G75-K104 interaction, we evaluated the effect of secondary mutation at G75 site on oncogenic K-Ras. Here, we substituted Glycine at 75 to Alanine (G75A) and measured transforming activity of each K-Ras mutant in Ba/F3 cells. Ba/F3 cells proliferate in an interleukin-3 (IL-3) dependent manner, but can be made IL-3 independent by exogenous KRAS expression [26]. Ba/F3 cells expressing K-Ras^G12D^ exhibited exponential cell growth in absence of IL-3, while cells expressing wild-type K-Ras or K-Ras^G12D;C185Y^, an inactive mutant that cannot be farnesylated, fail to survive [27]. Consistent with our findings *in vivo* K104Q mutation in combination with G12D decreases IL-3 independence. However, G75A in combination with G12D reduced growth Ba/F3 cells to a greater extent than K104Q (Fig. 4A). In line with this data, G75A mutant has a greater reduction on K-Ras activity, leading to decreased MAPK signaling activation relative to K104Q (Fig. 4B and 4C). These data demonstrate that G75A has a greater inhibitory effect on oncogenic K-Ras activity than K104Q.

**Figure 4.**
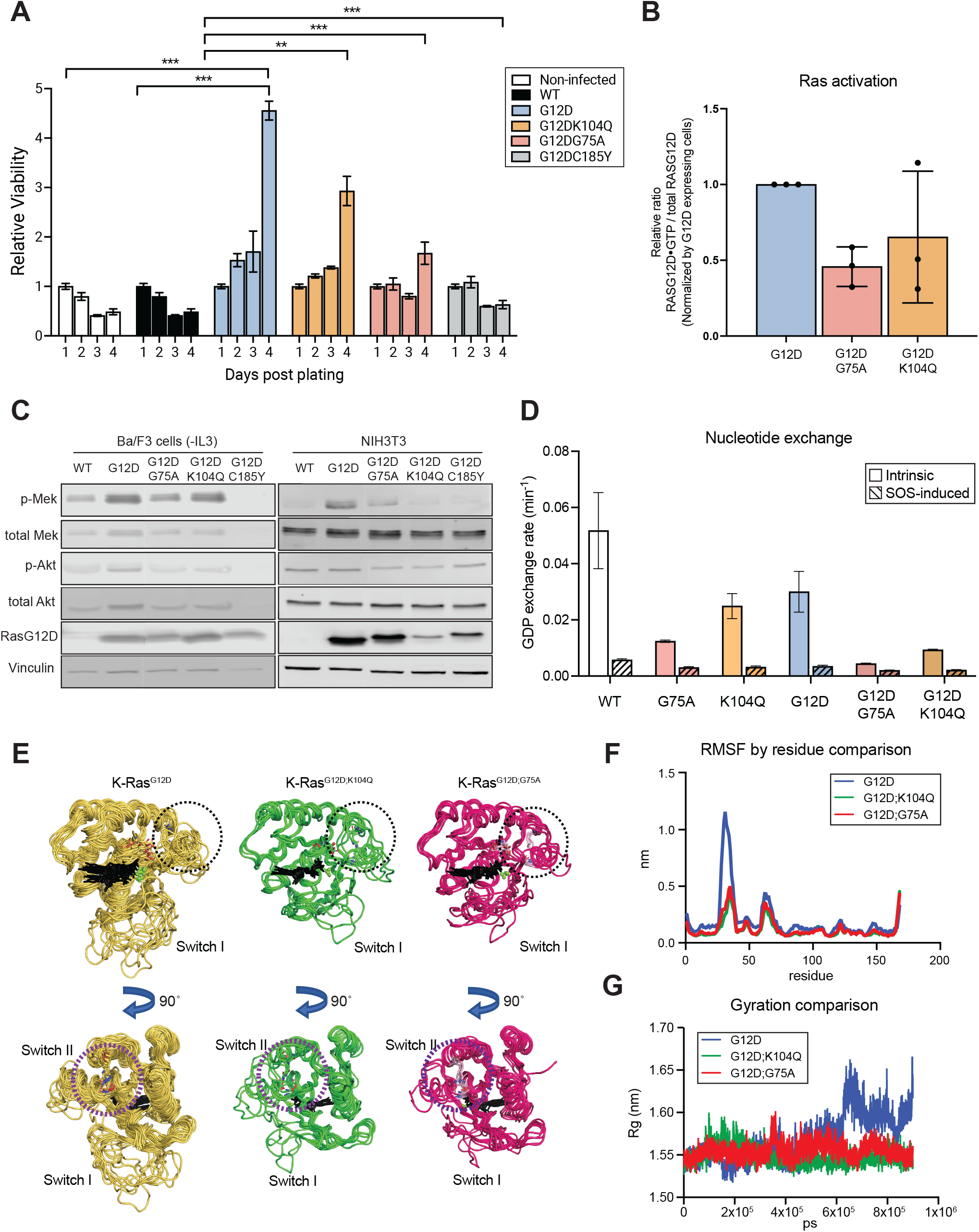
Modification at residues in allosteric regulatory network has a negative regulatory effect on oncogenic K-Ras. (A) The effect of secondary mutations at allosteric regulatory network on oncogenic K-Ras activity. Graph illustrated the cell viability of cells expressing the indicated K-Ras mutants. Oncogenic activity of K-Ras was measured using IL-3 dependency assay in Ba/F3 cells. For this assay, cells expressing the indicated K-Ras mutants were expanded in IL-3 containing media and plated into 96 well plate with IL-3 depleting media. The cell viability of each K-Ras mutant expressing cells was measured at day 1, 2, 3 and 4 post-plating. The cell viability of non-infected cells is gradually decreased in IL-3 depleted media, but oncogenic K-Ras expression rescues this IL-3 dependent phenotype. Secondary mutation at K104 or G75 suppresses oncogenic function of K-Ras. K-Ras^G12D;C185Y^, a loss of function mutant, was used as a negative control. Error bars represent SD. All measurements were normalized to cells of the same condition at day 1. ** P < 0.002, *** P < 0.005, **** P < 0.0001, based on a linear regression model. (B) Effect of secondary mutation at allosteric regulatory sites on oncogenic K-Ras activity. It was measured using Raf-RBD pulldown assay in NIH3T3 cells expressing the indicated K-Ras mutants. All measurement were normalized to cells expressing K-Ras^G12D^ at day 1. Each dot represents the measurement from an independent experiment. (C) Ras downstream signaling pathways analysis in cells expressing different K-Ras mutants. The activation of downstream signaling was assessed by Mek and Akt phosphorylation using western blotting for phosphorylation of Ser217/221 and Ser473 respectively. Ba/F3 cells expressing ectopic K-Ras mutants were grown in IL3 depleting media for 3 days prior to harvesting for western blotting. (D) Bar graph showing intrinsic and SOS-mediated nucleotide exchange rate for wild-type and single or double K-Ras mutants (K-Ras^G75A^, K-Ras^K104Q^, KRAS^G12D^, K-Ras^G12D;G75A^, and K-Ras^G12D;K104Q^). The rate of GDP release from mant-GDP-loaded KRAS protein was determined by monitoring the decrease in mant-GDP fluorescence signal as a function of time in the absence (intrinsic) and presence of SOS1. (E) Molecular dynamics simulations (900ns) for cluster analysis of GDP-bound K-Ras^G12D^, K-Ras^G12D;K104Q^ and K-Ras^G12D;G75A^. The black dotted and purple dashed circles show the position of switch II in the upper panels or the K104 allosteric site in lower panel, respectively. Sticks indicate the location of residues 12, 75 and 104. In the presence of co-mutation with G12D, switch II clusters between two conformations and switch I show a restricted range of dynamics. (F) Radius of gyration of K-Ras^G12D^, K-Ras^G12D;K104Q^ and K-Ras^G12D;G75A^ during 900ns MD simulation. (G) Root Mean Square Fluctuations (RMSF) of protein backbone during 900ns MD simulation. In K-Ras^G12D^, SW I shows the greatest dynamic range, however dynamic range at switch I is reduced in double mutant K-Ras.

To determine whether G75A mutant regulates oncogenic K-Ras activity through a similar mechanism as K104Q on SWII, we measured the effect of G75A on intrinsic and GEF-mediated nucleotide exchange of K-Ras. Consistent with *in vitro* data, G75A decreases both intrinsic- and GEF mediated nucleotide exchange, and it does so cooperatively with G12D (Fig. 4D). Cooperation of G12D with secondary mutation at G75 appeared to be greater than ones with at K104, suggesting that the G75-K104 interaction is critical for SWII interaction with SOS and K-Ras activation.

To determine if G75A mutation has a structurally similar effect on K-Ras as K104Q, we performed molecular dynamic simulations of K-Ras^G12D^, K-Ras^G12D;K104Q^, and K-Ras^G12D;G75A^. MD simulations demonstrated that both K-Ras^G12D;K104Q^ and K-Ras^G12D;G75A^ decreased dynamics in SWI and SWII compared to K-Ras^G12D^, resulting in a more compact structure (Fig. 4E-G). K-Ras^G12D^ progressively opens during simulation, resulting in dramatic opening of the active site (i.e. switch I and II), whereas additional mutations of K104 or G75 reduced opening of the active site (Fig. 4F and 4G). The effect of SWII on SWI in our simulations provides additional details into how these mutants suppress nucleotide exchange of K-Ras^G12D^. SOS catalyzing nucleotide exchange by stabilizing opening of the active site of K-Ras, and it does so by inserting a helix into the active site of K-Ras. It stands to reason that stabilization of the active site (i.e. SWI) in a more closed conformation would reduce the favorability of the K-Ras-SOS interaction. Thus, K104Q and G75A attenuate SOS exchange by regulating the conformation and dynamics of both SWI and SWII.

## Discussion

Overcoming mutational activation of K-Ras is a major hurdle in oncology today. KRAS mutations are common in cancers with high mortalities and are associated with poor clinical outcomes [28]. As a result, K-Ras is a high-value therapeutic target. Although there are many available small molecule inhibitors that target kinases downstream of K-Ras, effector-based therapy is complicated by the highly context-dependent nature of K-Ras signaling [18, 29]. By extension, the therapeutic approaches most likely to be broadly effective is to regulate the activity of K-Ras directly. Nevertheless, the Ras proteins are notoriously difficult to target due to their high affinity for GTP and the lack of accessible pockets for allosteric inhibitors [30]. In this study, we used genetics and structural biology to evaluate the therapeutic potential of targeting an allosteric regulatory network that controls nucleotide exchange for K-Ras.

Previously, we identified a post-translational modification - acetylation of lysine 104 - representing a unique allosteric regulatory site on K-Ras and demonstrated that K104Q mutation suppressed K-Ras^G12V^ oncogenicity in *in vitro* transforming assays [10]. We substituted lysine for glutamine to mimic constitutive acetylation of K104, however, it is not entirely clear how well this mutation really phenocopies actual acetylation. For example, consistent with our prior and current work, multiple groups demonstrated that the K104Q mutation interrupts interaction with GEF leading to reduction of GEF-induced nucleotide exchange, but other study found that acetylation of K104 did not [13, 31, 32]. Our modeled structure of K-Ras with acetylated K104 suggests that acetylation of K104 should reduce GEF-induced nucleotide exchange in K-Ras. Despite of this uncertainty, in this paper, we used the K104Q mutation to evaluate the function of perturbing SWII stability.

To determine if/how K104Q affects K-Ras function in a physiological setting, we generated genetically engineered mice in which the mutation is inserted into the endogenous *Kras* locus. In the context of wild-type K-Ras, we found that K104Q was not a strong loss-of-function mutation since animals expressing just a single allele of *Kras*^*K104Q*^ arose at Mendelian frequency and had no discernible phenotype (Table 1). This observation is consistent with a previous study in which ectopic K-Ras^K104Q^ restored proliferative capacity to mouse embryonic fibroblasts (MEFs) that lack all Ras proteins [13]. One remaining unknown is how little K-Ras activity is sufficient to support development *in vivo* or to promote proliferation in Rasless MEFs. In contrast to its indiscernible effect on wild-type K-Ras, K104Q reduces hyper-proliferative phenotype in the colonic epithelium through regulating MAPK signaling pathway by decreasing the oncogenic activity of K-Ras^G12D^ directly (Fig. 1). In addition, it rescues the differentiation phenotype in the small intestinal epithelium and increases survival in animals with colonic tumors (Fig. 1 and 2). These results clearly demonstrate the negative regulatory effect of K104Q on oncogenic K-Ras, thus our *in vivo* analysis is consistent with our previous *in vitro* findings.

**Table 1.** Expected and observed frequencies of offspring from crossing *Kras*^*K104Q/+*^ and *Kras*_LSL-G12D;K104Q/*+*_ mice.

| Kras Genotypes | Expected Frequency | Observed Frequency |
| --- | --- | --- |
| +/+ | 25% (21/84) | 27.4% (23/84) |
| LSL-G12DK104Q/+ | 25% (21/84) | 23.8% (20/84) |
| K104Q/+ | 25% (21/84) | 21.4% (18/84) |
| K104Q/LSL-G12DK104Q | 25% (21/84) | 26.2% (22/84) |

| *Kras* Genotypes | Expected Frequency | Observed Frequency |
| --- | --- | --- |
| +/+ | 25% (21/84) | 27.4% (23/84) |
| LSL-G12D;K104Q/+ | 25% (21/84) | 23.8% (20/84) |
| K104Q/+ | 25% (21/84) | 21.4% (18/84) |
| K104Q/LSL-G12D;K104Q | 25% (21/84) | 26.2% (22/84) |

With crystal structure analysis, we demonstrated that K104Q alters the conformation of the α2 helix of the SWII domain, which is primarily responsible for Ras-GEF interaction (Fig 3C and 3D). Consistent with this structural finding, K104Q reduced the rate of nucleotide exchange of K-Ras (Fig. 3B). While many of the common K-Ras activating mutations (e.g., G12D) reduce GAP-induced hydrolysis activity, their GEF-mediated nucleotide exchange activity is retained [33]. Moreover, it is likely that nucleotide exchange is required for full K-Ras activation because it can counter the intrinsic hydrolysis activity of many K-Ras mutants. Indeed, inhibition of GEF-induced nucleotide exchange was the proposed mechanism by which K104Q inhibited the *in vitro* transforming activity of K-Ras^G12V^ [10], and it is the mechanism through which covalent G12C inhibitors are proposed to inhibit oncogenic K-Ras function [8]. Intriguingly, oncogenic G12 mutant causes a shift in the population of local conformational state at SWII and α3 helix, leading to alteration in SWII flexibility, while wild-type K-Ras does not (Fig. 3C and 3D) [34, 35]. This conformational change at SWII caused by G12 mutation cooperates with K104Q, leading to greater reduction of GEF-induced nucleotide exchange nearly to the rate of intrinsic exchange (Fig. 3B). This elucidates how K104Q preferentially affects the function of K-Ras^G12D^ over WT K-Ras, as suggested by our *in vivo* data.

Our structural analysis revealed that K104 plays a key role in stabilizing the SWII alpha helix through its interaction with M72, R73 and G75, and destabilization of SWII in the K104Q mutant is likely due to the loss of interaction between K104 and R73 (Fig. 3D and 3F). This is consistent with the previous NMR analysis of K-Ras^K104Q^ mutant, which indicated that K104Q mutant only perturbs protein conformation proximal to the site of mutation in helix α3 as well as the end of helix α2 in SWII [13]. Furthermore, modeling of Ac-K104 within this network suggests a greater perturbation - loss of interaction between K104 and R73 and M72 - which is predicted to further destabilize SWII. Interestingly, both modifications at K104 site retain the interaction with G75, suggesting that K104-G75 interaction in allosteric network might be primary for keeping stabilization of SWII (Fig. 3F). In line with this, G75A abolishes nucleotide exchange of K-Ras^G12D^ (Fig. 4D). Moreover, it has a stronger negative regulatory effect on oncogenic activity of K-Ras than K104Q (Fig. 4A and 4B).G75A and K104Q seem to have a similar molecular mechanism on modulating oncogenic K-Ras activity, as suggested by our MD simulation (Fig. 4E - 4G). One remaining question is that how G75A has a stronger impact than K104Q mutant. One possibility is losing interaction with M72. In our MD simulation, G75A completely lost interaction with M72 while K104Q retained (Data not shown). These data suggest that Ac-K104 might have a greater impact on oncogenic activity of K-Ras^G12D^ than K104Q because of similar predicted model of Ac-K104 and G75 in allosteric network. However, this question remains to be determined further.

In light of our data from biochemical assays and computational stimulation, K104Q or G75A perturb allosteric network at SWII but not completely destroy. These mutations cause conformational changes and reduced dynamics at SWII and SWI domains leading to affecting nucleotide exchange primarily, but not binding affinity to downstream effectors (Fig. 3B, 4D and S5B). Considering K104Q is not a loss of function mutation, these suggest that the level of perturbing is important to keep adequate level of K-Ras activity. Interestingly, G75A has been reported as a single nucleotide polymorphism that occurs at a frequency of 4×10^−6^ in human. Although rare, it is interesting to consider that individuals carrying the G75A might be resistant to *KRAS* activating mutations that occur on the allele. Including G75A, we found 32 other germline SNPs in human *KRAS* gene (Table S2) and mapped the location of identified SNPs in K-Ras structure. Among these germline SNPs, only few SNPs including G75A are located near SWII, reported as a part of proposed inhibitor binding pocket (Fig. S6) [36]. Addition to clinical benefit prediction, germline SNPs provides valuable information to validate loss-of-function mutation for developing therapeutic approaches to target K-Ras mutant. In conclusion, our data suggest the perturbing allosteric network in SWII might be a promising therapeutic approach to target oncogenic K-Ras mutant in human cancers harboring *KRAS* mutations.

## Materials and Methods

### Generating *Kras*^*LSL-G12D;K104Q*^ and *Kras*^*K10Q*^ mouse model using CRISPR/Cas9 system

The *Kras*^*LSL-G12D;K104Q*^ and *Kras*^*K10Q*^ allele were generated in J1 embryonic stem (ES) cells (129S4/SvJae genetic background) with *Kras*^*LSL-G12D*^ allele in which the G12D point mutation is targeted in one of endogenous K-Ras locus following a transcriptional stop element (lox-stop-lox, LSL) [14]. We engineered a lysine to glutamine mutation at codon 104 in exon 4 to generate *Kras*^*LSL-G12D;K104Q*^ allele using CRISPR/Cas9. Briefly, the guide RNA sequence against the genomic sequence in exon4 of the K-Ras alleles was cloned into px459 plasmid as described in [37]. The single strand deoxynucleotides (ssODN) repair template containing single base pair changes to generate point mutation at codon 104 and a silent mutation to create NcoI restriction enzyme site for genotyping is used for generating these mouse models (Table S3). *Kras*^*LSL-G12D*^ ES cells were electroporated using the Lonza Nucleofector system according to manufacturer’s instruction. Homozygote ES cell clone was utilized for generating both *Kras*^*LSL-G12D;K104Q*^ and *Kras*^*K10Q*^ mouse model. Positive ES clone was injected into C57BL/6J embryos in the Beth Israel Medical Center Transgenic Core and chimeric males were backcrossed to C57BL/6 females.

### Ras activation assay and immunoblotting

Glutathione *S*-transferase protein fused Raf binding domain (GST-RBD) conjugated beads were produced as described [38]. Frozen tissue or cell pellets were lysed in MLB lysis buffer [25mM HEPES, 150mM NaCl, 1% NP-40, 0.25% Sodium Deoxycholate, 10% glycerol and 10mM MgCl2] supplemented with cOmplete™ protease inhibitor cocktail (Roche), and phosphatase inhibitor cocktail 2 and 3 (Sigma Aldrich). Equal amount of protein lysates was incubated with 20ug of GST-RBD beads. The precipitated proteins were resolved on a 12% SDS-PAGE gel, and immunoblotting was performed to detected GTP-bound Ras proteins using anti-K-Ras antibody (Proteintech, 12063-1-AP), anti-RasG12D mutant specific antibody (Cell Signaling Technologies [CST], 14429) or anti-Ras antibody (Millipore, 05-516). To analyze Ras downstream signaling components, immunoblotting was performed to standard protocols. Immunoblot images were quantified using Image Studio Software (LI-COR_®_). Primary antibodies included: anti-Ras G12D (CST, 14429), anti-Vinculin (Sigma Aldrich, V9131), anti-pErk1/2(Thr202/Tyr204; CST, 4377), anti-Erk1/2 (CST, 4696), anti-Mek (Ser217/221; CST, 9121), anti-Mek (CST, 4694), anti-p-Akt (Ser473; CST, 4060), and anti-Akt (CST, 9272). Secondary antibodies included: anti-mouse IgG Alexa Fluor 680 (Invitrogen, A21058) and anti-rabbit IgG Alexa Fluor 800 (Invitrogen, A32735).

### Tissue staining, IHC, and IF

Tissue sections (5μm) were deparaffinized in a standard xylene and ethanol series and stained according to the Standard H&E staining protocols. For colonic crypt height analysis, colonic crypts were measured using Olympus VS-ASW version 2.7 software as previously described [39]. Immunohistochemistry (IHC) and immunofluorescence (IF) were performed as previously described [18]. IHC Images were acquired using an Olympus BX-UCB slide scanner and IF images were acquired using a Zeiss LSM 880 upright system. Primary antibodies included: anti-pH3 (Ser10; CST, 9701), anti-E-Cadherin (BD bioscience, BDB610181), anti-β-catenin (CST, 9561), and anti-lysozyme (Thermo Scientific, RB-372-A1). Fluorescent secondary antibodies included: anti-mouse IgG2a-Alexa Fluor 488 (Invitrogen, A11034) and anti-rabbit Alexa Fluor 594 (Invitrogen, A21125). Translocation of β-catenin on colon tumor sections were analyzed by QuPath [40].

### Animal studies

Animal studies were approved by the Institutional Care and Use Committee at Beth Israel Deaconess Medical Center. Mice were fed ad libitum and housed in a barrier facility with a temperature-controlled environment and twelve-hour light/dark cycle. *Fabp1-Cre* (Strain 01XD8), *Apc*^*2lox14*^ (Strain 01XP3), and *Kras*^*LSL-G12D*^ (Strain 01XJ6) mice were obtained from the NCI Mouse Repository. Experimental animals were maintained on genetic background that was 80-95% C57BL/6. For colonic epithelium analysis, all animals were sacrificed at 8-12 weeks of age.

### Small molecule inhibitor treatments in murine colonic organoids

Mouse colonic organoids were maintained in maintained in advanced DMEM/F-12 supplemented with 1X Primocin (Invivogen #ant-pm-1), 10mM HEPES, 2mM Glutamax (Thermo Fisher #35050061), 500μM N-acetylcysteine, 25ng/ml EGF (Gibco, #PHG0311), and 65% WRN conditional medium produced as previously described [41]. For inhibitor treatment, organoids were dissociated into single cells and plated in 384 well at 1×10^3^ cell concentration. 24 hours post plating, compounds were added to each well over 12-point dose curves along with DMSO controls using a D300e digital drug printer (Tecan Life Sciences). The cell viability was assessed at day 6 post inhibitor treatment using CellTiter Glo 3D (Promega) according to manufacturer’s instruction.

### Cloning, expression, and purification of recombinant proteins

DNA constructs for the expression of human K-RAS 4b(1-169), K-RAS 4b(1-169)^G12D^, K-RAS 4b(1-169)^K104Q^, K-RAS 4b(1-169)^G12D;K104Q^, K-RAS 4b(1-169)^G75A^, K-RAS 4b(1-169)^G12D;G75A^, SOS^cat^(564-1048), RAF1(52-131), PI3K and NF1(1198-1530) in the format of His6-MBP-tev-POI were generated as described previously [42]. All expressed proteins were purified as outlined for KRAS4b(1-169) previously [43]. Briefly, the expressed proteins were purified from clarified lysates by IMAC, treated with His6-TEV protease to release the target protein, and the target protein separated from other components of the TEV protease reaction by a second round of IMAC. Proteins were further purified by gel-filtration chromatography in buffer containing 20 mM HEPES, pH 7.3, 150 mM NaCl, 2 mM MgCl2 (KRAS only), and 1 mM TCEP. The peak fractions containing pure protein were pooled, flash-frozen in liquid nitrogen, and stored at -80ºC.

### *in vitro* nucleotide exchange assay

Nucleotide exchange rate was measured by a fluorescence-based assay using mant-GDP as described previously [44]. For this assay, 2.5 µM K-Ras-mant-GDP was diluted into 20 mM HEPES pH 7.4, 150 mM NaCl, 5 mM MgCl_2_, and 1 mM TCEP in a 3 ml cuvette. The reaction was initiated by the addition of 3.5 mM GDP in the presence and absence of 1.5 µM SOS_cat_. The change in fluorescence signal was recorded using an excitation and emission wavelength of 355nm and 448nm every 15 seconds. Fluorescence traces were fitted into single exponential dissociative function using Mathematica’s nonlinear regression analysis.

### *in vitro* GTP hydrolysis assay

GTP hydrolysis rate was measured using a fluorescence-based assay, as previously described [45]. For this assay, K-Ras proteins (3 µM final concentration) were diluted into 50 mM Tris-HCl pH 7.5, 1 mM DTT, and 1 mM MgCl_2_ containing 4.5 µM phosphate-binding protein labeled with a phosphate sensor (MDCC, Fisher Scientific) for intrinsic GTPase activity or containing 1 nM NF1-GAP for GAP mediated GTPase activity. Fluorescence signals were measured using excitation and emission wavelengths of 430nm and 550nm every 30 seconds in plate reader (Tecan M1000). Fluorescence traces were fitted into exponential associative function and the rate constants were extracted using Mathematica’s nonlinear regression analysis.

### ITC assay

To measure binding affinities of GDP-bound wild-type K-Ras, K-Ras^K104Q^, K-Ras^G75A^, K-Ras^G12D^, K-Ras^G12D;G75A^ and K-Ras^G12D;K104Q^ to Raf1-RBD or Pi3k-RBD, 60 μM of K-Ras WT or mutant proteins and 600 μM of Raf1-RBD or Pi3k-RBD were placed in the cell and syringe, respectively. ITC experiments were performed in a MicroCal PEAQ-ITC (Malvern) at 25 °C using 19 injections of 2.2 μl injected at 150-s intervals. Data analysis was performed based on a binding model containing “one set of sites” using a nonlinear least-squares algorithm incorporated in the MicroCal PEAQ-ITC analysis software (Malvern).

### Crystallization and data collection

Crystallization was carried out using the sitting-drop vapor diffusion method by mixing GDP-bound K-Ras mutant proteins (10-15 mg/ml) with an equal volume of reservoir solution. Initial crystallization hits obtained from ammonium sulfate screens for both K-Ras^K104Q^ (0.1 M Tris pH 8.5 and 2 M ammonium sulfate) and K-Ras^G12D;K104Q^ (3.0 % Xylitol, 0.2 M ammonium acetate and 2.2 M ammonium sulfate) were further optimized to improve the diffraction quality. Optimized crystals were harvested for data collection and cryoprotected with a combination of 15% (v/v) PEG 3350 and 15% (v/v) glycerol solution before being flash-frozen in liquid nitrogen. Diffraction data sets were collected at 24-ID-C/E beamlines at the Advanced Photon Source (APS), Argonne National Laboratory. Crystallographic datasets were integrated and scaled using the XDS [46]. The crystal parameters and the data collection statistics are summarized in Table S1.

### Structure determination and analysis

K-Ras structures were solved by molecular replacement using the program Phaser as implemented in the Phenix/CCP4 suite of programs with PDB entry 5US4 (GDP-bound KRas^G12D^) [47-49]. Crystals for both structures exhibited twinning and pseudo-translational pathologies as diagnosed by the program Xtriage within the Phenix suite. The initial solution obtained from molecular replacement was refined using twin operator *-h, -k, l* in Refmac5 of the CCP4 suite, and the resulting *Fo-Fc* map showed clear electron density for the GDP nucleotide. The model was further improved using iterative cycles of manual model building in COOT [50] and refinement using Refmac5. The GDP nucleotide was placed in the nucleotide-binding pocket and followed by the addition of solvent molecules by the automatic water-picking algorithm in COOT. These water molecules were manually checked during model building. Alternative conformations of amino acid side chains were added using COOT during final rounds of refinements. The refinement statistics for the structures are summarized in Table S1. All the figures were rendered in PyMOL (Schrödinger, LLC). Crystallographic and structural analysis software support is provided by the SBGrid consortium [51].

### Data deposition

The atomic coordinates and structure factors of GDP-bound K-Ras^K104Q^ and K-Ras^G12D;K104Q^ reported in this paper have been deposited in the Protein Data Bank and were assigned accession codes 6WS2 and 6WS4, respectively.

### Plasmid construction, cell lines

pLx307 EGFP and pLX307 K-Ras^G12D^ plasmids, and Ba/F3 cells were kindly gifted from Andrew Aguirre (Dana-Farber Cancer Institution). All K-Ras secondary mutant and wild-type K-Ras constructs were generated by a PCR based strategy using a site-directed mutagenesis kit (Stratagene) according to the manufacturer’s instructions. HEK293T and NIH3T3 cells were obtained from the American Type Culture Collection (ATCC; Manassas, VA). All cells were cultured in recommended media at 37°C, 5% CO2. Lentivirus was produced as previously described [10].

### Functional assay for determining oncogenic K-Ras activity in vitro

To measure the regulatory effect of secondary mutants on K-Ras^G12D^ *in vitro*, Ba/F3 cells stably expressing different K-Ras mutant were plated in 96 well plate at 1×10^3^ cells concentration with IL-3 depleting media. The cell viability was assessed using CellTiter-Glo 2.0 assay (Promega) according to the manufacturer’s instruction.

### MD simulations

Starting coordinates for molecular dynamic simulation were generated from the starting crystal structure of wildtype GDP-bound K-Ras (PDB code 6MBU) [43]. Refined structure was then mutated *in silico* and used for the online platform ‘solution builder’ available from CHARM-GUI to prepare starting files for energy minimization and molecular dynamics simulation [52-54]. Charged residues, including protein termini, were protonated, or deprotonated in accordance with neutral pH. A cubic box with edges 10 Å from each protein was created and filled with TIP3P water molecules and neutralized with Cl^-^ and Na^+^ ions to 150 mM. Minimizations, equilibrations, and simulations were done using GROMACS (ver. 2020.1) and a GPU server featuring 8x Tesla v100 workstation, on the O2 High Performance Compute Cluster, supported by the Research Computing Group, at Harvard Medical School. Solvated systems were energy minimized by steep integration for 5000 steps or at a maximum force of 1000 kJ/mol/nm or less. The Verlet cutoff scheme was used for nonbonded atoms and the LINCS algorithm was applied to covalent H-bonds. Short-range van der Waals interactions were switched off from 1.0-1.2 nm, and long-range interactions were computed using the Particle Mesh Ewald method. Finally, simulations were performed using the GROMOS force-field. Validation and analyses were done in GROMACS, including radius of gyration, RMSF calculation, and cluster analysis using the GROMOS algorithm [55]. Distance measurements and visual analyses were done using PyMOL and VMD [56].

## Authors’ Contributions

**Conception and design**: M. Yang, D. Simanshu, and K. Haigis

**Acquisition of Data**: M. Yang, T. Tran, B. Hunt, R. Agnor, T. Waybright, C. Johnson and J. Nowak,

**Data analysis**: M. Yang, T. Tran., C. Johnson and D. Simanshu

**Writing, review, and/or revision of manuscript**: M. Yang, T. Tran, D. Simanshu, and K. Haigis

**Study supervision**: D. Simanshu and K. Haigis

## Acknowledgments

We thank Dominic Esposito, William Gillette, Jennifer Mehalko, Vanessa Wall, John-Paul Denson, Jose Sanchez Hernandez, Nitya Ramakrishnan, Troy Taylor, Allison Champagne, Simon Messing, Peter Frank, Min Hong, Matt Drew and Kelly Snead of the FNLCR Protein Expression Laboratory for their help in cloning, expression and purification of recombinant proteins. We are thankful to Andrew Stephen and Srisathiyanarayanan Dharmaiah for their help with biochemical assays. We are grateful to the staff of 24-ID-C/E beamlines at the Advanced Photon Source, Argonne National Laboratory, for their help with data collection. This work was supported by NCI grant (R01-CA178017) to K.M.H. and funded in part with federal funds from National Cancer Institute, NIH Contract HHSN261200800001E. Part of this work is based on research conducted at the Northeastern Collaborative Access Team beamlines, which are funded by the National Institute of General Medical Sciences from National Institutes of Health Grant P41 GM103403. This research used resources of the Advanced Photon Source, a US Department of Energy (DOE) Office of Science User Facility operated for the DOE Office of Science by Argonne National Laboratory under Contract DE-AC02-06CH11357. The content of this publication does not necessarily reflect the views or policies of the Department of Health and Human Services, and the mention of trade names, commercial products, or organizations does not imply endorsement by the US Government.

