## Supplementary Figure 1 for "Allosteric regulation of switch-II controls K-Ras oncogenicity"

### Figure S1

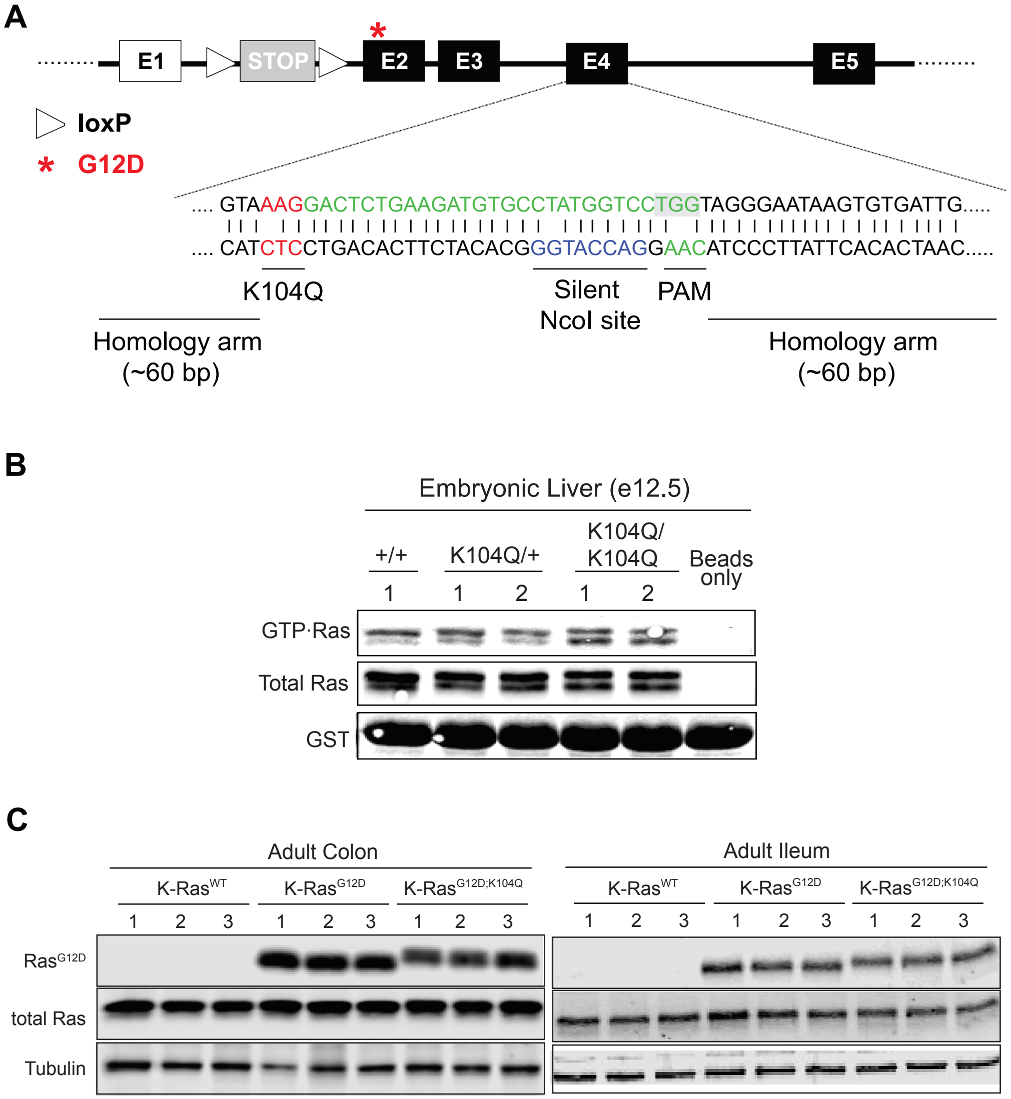


**Figure S1. Generation of K-Ras$^{LSL-G12D;K104Q}$ and K-Ras$^{K104Q}$ mouse model.** (A) Schematic illustrating the CRISPR/Cas9 design to generate the K104Q mutation in Kras$^{LSL-G12D/+}$ ES cells. The K104Q mutation was inserted into both Kras alleles. (B) Western blotting analysis for Ras activation in embryonic livers carrying indicated the *Kras* alleles. The level of Ras-GTP was determined by Raf-RBD pulldown assay. Each lane contains lysate of embryonic liver from an individual embryo. (C) Endogenous Ras expression in the adult small and large intestines. Each lane contains intestine lysate from an individual Fabp1-Cre animal carrying various Kras genotype. Total Ras levels were detected with a pan-Ras antibody and RasG12D was detected with a mutant-specific
