## Supplementary Figure 2 for "Allosteric regulation of switch-II controls K-Ras oncogenicity"

**Figure S2**

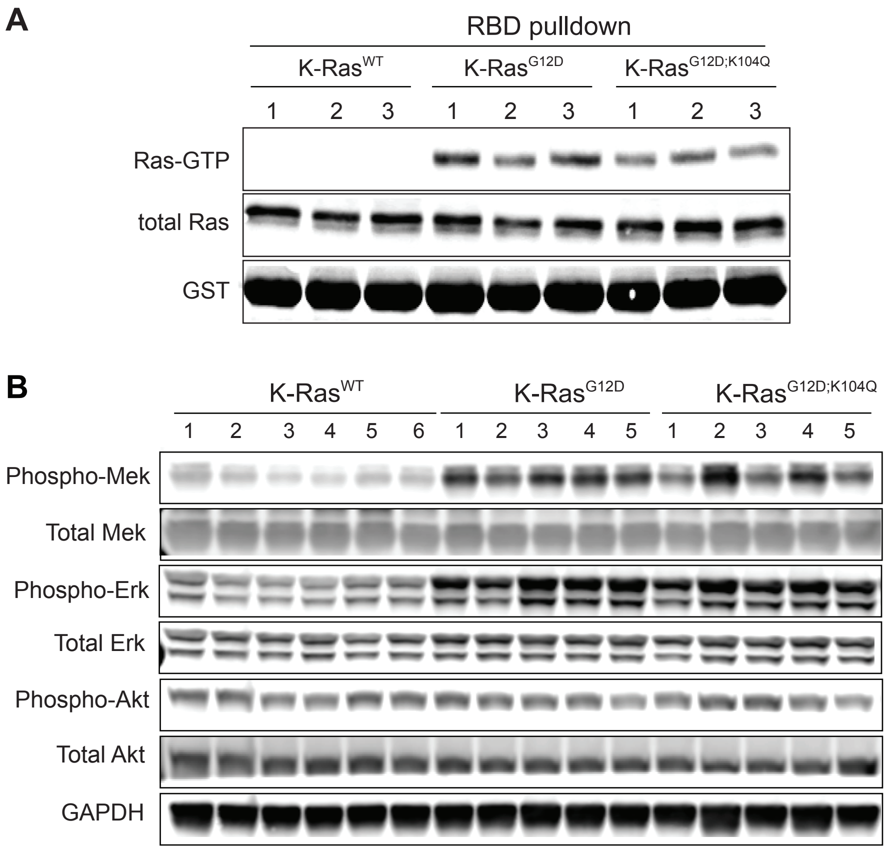


**Figure S2. Analysis of colons from *Fabp1-Cre* mice expressing different K-Ras mutants.** (A) Western blot analysis for Ras activation. The level of Ras-GTP was measured using an Raf-RBD pulldown assay. Each lane contains lysate from an individual mouse. (B) Western blotting for Ras downstream signaling components in colons carrying the indicated *Kras* alleles. Each lane contains lysate from an individual mouse.
