## Supplementary Figure 3 for "Allosteric regulation of switch-II controls K-Ras oncogenicity"

### Figure S3

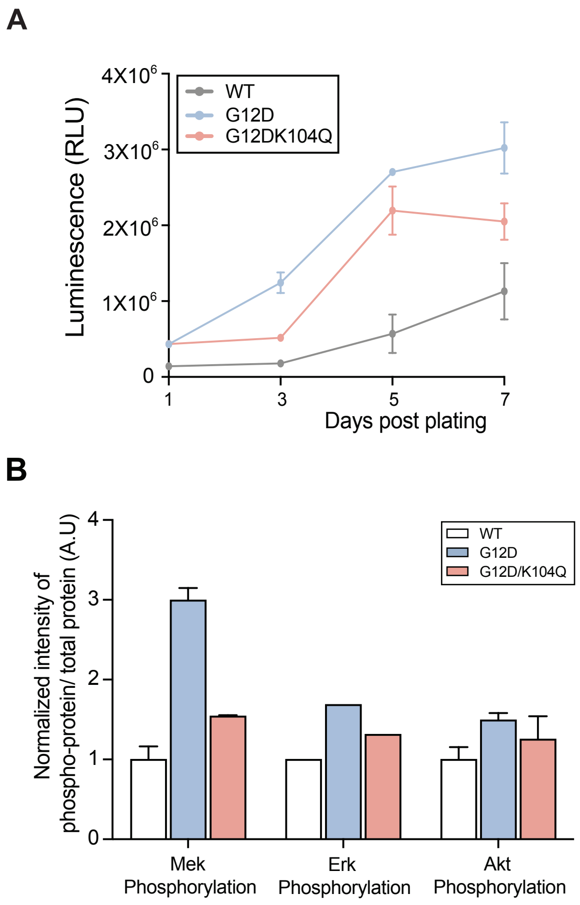


**Figure S3. the effect of K104Q mutant on oncogenic K-Ras$^{G12D}$ in mouse colonic epithelium organoids system.** (A) Proliferation analysis of colonic organoids expressing different K-Ras mutants. The proliferation of organoids was measured at day 1, 3, 5 and 7 post plating using Celltiter Glo 3D assay. The K104Q mutation attenuates enhanced proliferation of K-Ras$^{G12D}$ organoids. (B) Activation of Ras downstream signaling pathways in colonic organoids carrying various Kras alleles. The K104Q mutant represses elevated MAPK signaling activation in colonic organoids expressing K-Ras$^{G12D}$. Ras downstream signaling activation was assessed by Mek, Erk and Akt phosphorylation using western blotting for phosphorylation of Ser217/221, Thr202/204 and Ser473 respectively.
