## Supplementary Figure 4 for "Allosteric regulation of switch-II controls K-Ras oncogenicity"

### Figure S4

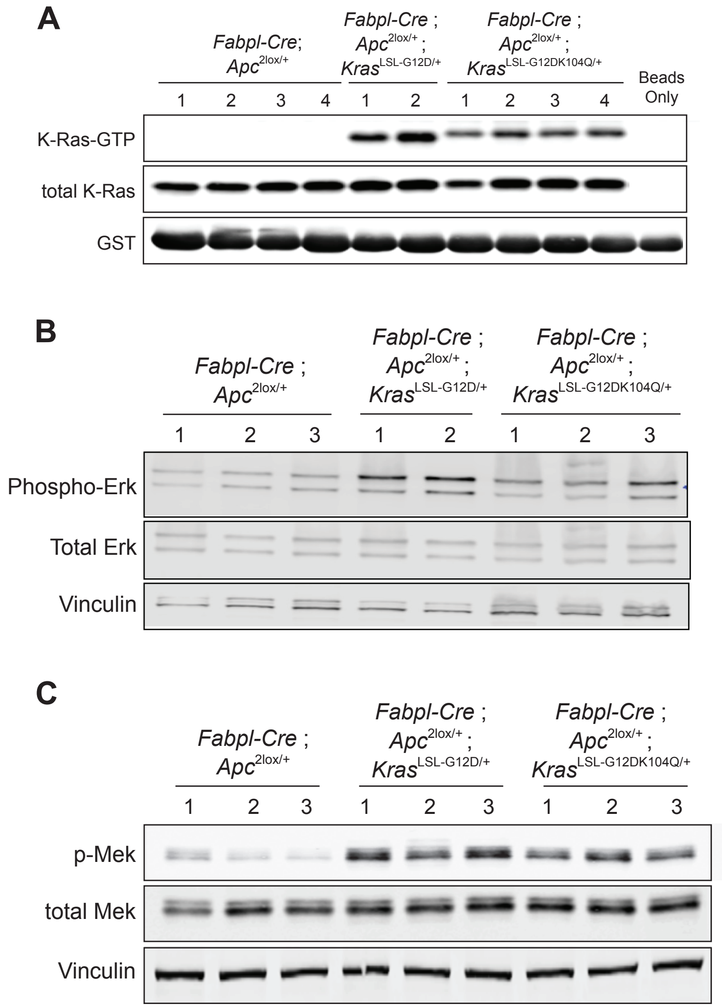


**Figure S4. Analysis of Ras downstream pathway activation in mouse colon tumors.** (A) Western blotting analysis for assessing Ras activation using Raf-RBD pulldown analysis in colon tumors of the noted genotypes. (B-C) western blotting analysis for determining the activation of Ras downstream signaling effectors, Erk (B) and Mek (C) in colon tumors from indicated genotypes. In all figures, each lane contains lysate from tumors harvested from an individual mouse.
