## Supplementary Figure 5 for "Allosteric regulation of switch-II controls K-Ras oncogenicity"

### Figure S5

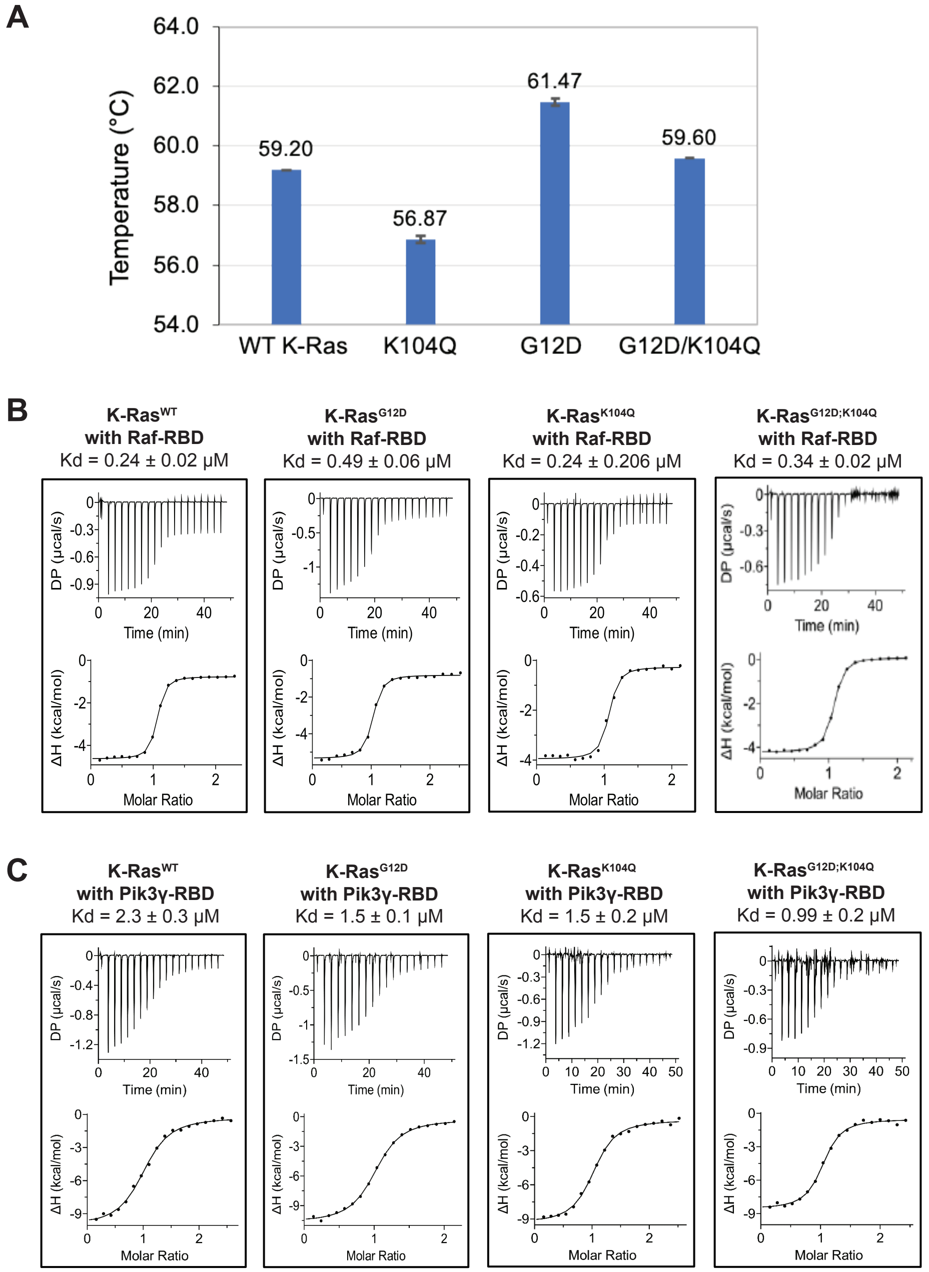


**Figure S5. Effect of the K104Q mutation on K-Ras thermal stability and its interaction with downstream effectors.** (A) Bar graph showing the melting temperature (Tm) of GDP-bound forms of K-Ras. Results are plotted as the mean ± S.D. (N = 3). (B-C) ITC titration experiments to measure the dissociation constant of Raf-RBD (B) or Pik3γ-RBD (C) with GDP bound K-Ras. The DP is measured differential power between the reference cells and the sample cells.
