## Supplementary Figure 6 for "Allosteric regulation of switch-II controls K-Ras oncogenicity"

### Figure S6

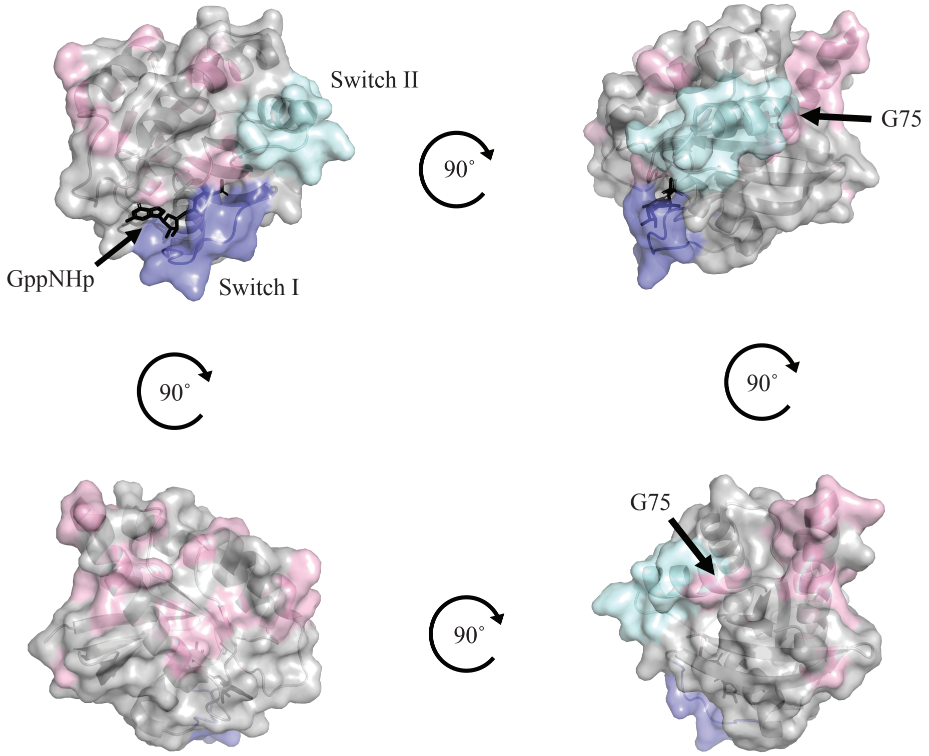


**Figure S6. Depiction of human germline codon substitutions in K-RAS**. Location of amino acid changes as a result of germline SNPs are shown in light pink on the surface of K-RAS (PDB code 6GOF). Switches I and II are labeled in the first panel and by purple and cyan surfaces, respectively. The location of G75 is labelled in the upper right panel for reference. Bound nucleotide is shown as black sticks. Germline data for SNPs resulting in a amino acid change were taken from gNOMAD database.
