## Supplementary Table 1 for "Allosteric regulation of switch-II controls K-Ras oncogenicity"

**Table S1. Crystallographic data collection and refinement statistics.**

| | GDP-bound KRAS-K104Q | GDP-bound KRAS G12D-K104Q |
|---|---|---|
| **Data collection** | | |
| Resolution range (Å) | 42.28 - 1.59 (1.65 - 1.59) | 42.47 – 1.84 (1.91 - 1.84) |
| Space group | P 3 | P 3 |
| Unit cell<br>a, b, c (Å)<br>α, β, γ (°) | 84.56, 84.56, 88.36<br>90, 90, 120 | 84.93, 84.93, 89.30<br>90, 90, 120 |
| Total reflections | 479681 (77977) | 427375 (70724) |
| Unique reflections | 94111 (15241) | 62031 (10046) |
| Multiplicity | 5.1 (5.1) | 6.9 (7.0) |
| Completeness (%) | 99.7 (99.8) | 100 (100) |
| Mean I/sigma(I) | 11.33 (2.01) | 10.09 (2.69) |
| Wilson B-factor ($Å^2$) | 22.49 | 27.03 |
| R-merge | 0.076 (0.717) | 0.119 (0.555) |
| R-meas | 0.085 (0.803) | 0.129 (0.602) |
| CC1/2 | 0.997 (0.775) | 0.994 (0.779) |
| **Refinement** | | |
| Reflections used in refinement | 94107 (9309) | 62001 (5680) |
| Reflections used for R-free | 4763 (505) | 2995 (288) |
| R-work | 0.1816 (0.3211) | 0.1333 (0.2620) |
| R-free | 0.1954 (0.3293) | 0.1580 (0.2942) |
| Number of non-H atoms | 6088 | 5777 |
| macromolecules | 5439 | 5409 |
| ligands | 152 | 132 |
| solvent | 497 | 236 |
| Protein residues | 676 | 676 |
| RMS - bond length (Å) | 0.0053 | 0.0085 |
| RMS - bond angle (°) | 1.37 | 1.56 |
| Ramachandran favored (%) | 97.90 | 98.05 |
| Ramachandran allowed (%) | 2.10 | 1.95 |
| Ramachandran outliers (%) | 0.00 | 0.00 |
| Average B-factor ($Å^2$) | 28.07 | 32.71 |
| macromolecules | 27.48 | 32.77 |
| ligands | 25.75 | 27.08 |
| solvent | 35.23 | 34.58 |

Note: Statistics for the highest-resolution shell are shown in parentheses.
