## Supplementary Table 2 for "Allosteric regulation of switch-II controls K-Ras oncogenicity"

**Table S2. List of missense germline variants in *Kras* allele.**

| Protein Consequence | rsID | Allele Frequency | Annotation | Source |
|---|---|---|---|---|
| p.Leu6Val | rs1296330213 | 3.18492E-05 | missense_variant | gnomAD Genomes |
| p.Thr50Ile | rs1407509439 | 3.18492E-05 | missense_variant | gnomAD Genomes |
| p.Gly60Val | rs727503108 | 3.97886E-06 | missense_variant | gnomAD Exomes |
| p.Thr74Ala | rs770020203 | 3.97937E-06 | missense_variant | gnomAD Exomes |
| p.Gly75Ala | rs780974222 | 3.9795E-06 | missense_variant | gnomAD Exomes |
| p.Gly77Ala | rs756890312 | 3.97991E-06 | missense_variant | gnomAD Exomes |
| p.Asp108Tyr | rs763553461 | 7.96844E-06 | missense_variant | gnomAD Exomes |
| p.Val112Ile | rs775836436 | 3.98283E-06 | missense_variant | gnomAD Exomes |
| p.Thr124Ser | rs575569675 | 3.9813E-06 | missense_variant | gnomAD Exomes |
| p.Asp126His | rs1363431968 | 3.98137E-06 | missense_variant | gnomAD Exomes |
| p.Thr127Arg | rs781634879 | 3.98162E-06 | missense_variant | gnomAD Exomes |
| p.Lys128Arg | rs746609817 | 3.98134E-06 | missense_variant | gnomAD Exomes |
| p.Ala130Thr | rs1463850736 | 3.9814E-06 | missense_variant | gnomAD Exomes |
| p.Ala130Ser | rs1463850736 | 3.1902E-05 | missense_variant | gnomAD Genomes |
| p.Ala130Val | rs730880473 | 1.9907E-05 | missense_variant | gnomAD Exomes |
| p.Ser136Asn | rs757816355 | 1.1944E-05 | missense_variant | gnomAD Exomes |
| p.Gly138Arg | rs778702415 | 3.98156E-06 | missense_variant | gnomAD Exomes |
| p.Asp154Gly | rs989151052 | 3.98181E-06 | missense_variant | gnomAD Exomes |
| p.Gly138Glu | rs754870563 | 3.98149E-06 | missense_variant | gnomAD Exomes |
| p.Ile142Thr | rs1344202459 | 3.18959E-05 | missense_variant | gnomAD Genomes |
| p.Thr158Ile | rs749177256 | 3.98178E-06 | missense_variant | gnomAD Exomes |
| p.Val160Met | rs755877953 | 2.12353E-05 | missense_variant | gnomAD Exomes, gnomAD Genomes |
| p.Ala155Gly | rs755177746 | 4.01629E-06 | missense_variant | gnomAD Exomes |
| p.Ile163Val | rs1470495974 | 3.99383E-06 | missense_variant | gnomAD Exomes |
| p.Arg164Gln | rs372793780 | 1.77673E-05 | missense_variant | gnomAD Exomes,gnomAD Genomes |
| p.Tyr166His | rs397517476 | 7.96216E-06 | missense_variant | gnomAD Exomes |
| p.Lys167Arg | rs1417039463 | 3.18492E-05 | missense_variant | gnomAD Genomes |
| p.Lys170Glu | rs1191739287 | 3.18512E-05 | missense_variant | gnomAD Genomes |
| p.Ile171Met | rs766231905 | 1.76949E-05 | missense_variant | gnomAD Exomes,gnomAD Genomes |
| p.Ser172Cys | rs772985440 | 3.98051E-06 | missense_variant | gnomAD Exomes |
| p.Glu174Lys | rs771629239 | 3.98057E-06 | missense_variant | gnomAD Exomes |
| p.Gly179Ser | rs200970347 | 0.00049184 | missense_variant | gnomAD Exomes,gnomAD Genomes |
| p.Lys179Arg | rs763736188 | 8.01321E-06 | missense_variant | gnomAD Exomes |
| p.Thr183Lys | rs1256144582 | 4.01184E-06 | missense_variant | gnomAD Exomes |
| p.Ile183Val | rs529925358 | 3.18494E-05 | missense_variant | gnomAD Exomes |
| p.Arg164Gln | rs758575947 | 1.9907E-05 | missense_variant | gnomAD Exomes |
| p.Cys185Tyr | rs775000854 | 4.01081E-06 | missense_variant | gnomAD Exomes |
| p.Val186Ile | rs1432921036 | 1.20349E-05 | missense_variant | gnomAD Exomes |
| p.Ile187Val | rs779951033 | 3.98143E-06 | missense_variant | gnomAD Exomes |
| p.Ile188Val | rs755967833 | 1.06181E-05 | missense_variant | gnomAD Exomes,gnomAD Genomes |
| p.Met188Ile | rs1336314580 | 4.01661E-06 | missense_variant | gnomAD Exomes |
| p.Met189Leu | rs201170656 | 0.000265427 | missense_variant | gnomAD Exomes,gnomAD Genomes |
