## Supplementary Table 3 for "Allosteric regulation of switch-II controls K-Ras oncogenicity"

**Table S3. Primers and oligonucleotides used for genotyping and CRISPR.**

| Name | Sequence |
|---|---|
| K104Q genotyping Forward primer | 5’- CCATCCCTAGTTTCAGGCACT-3’ |
| K104Q genotyping Reverse primer | 5’- CATGTATACCCCTAATGATGCCATCAGG-3’ |
| Repair template for K104Q mouse model | CTTATGCTTCTGCTTCACTTTGTTTCTTCCCCAGAGAACAAATTAAAAG AGTACAGGACTCTGAAGATGTGCCCATGGTCCTCGTAGGGAATAAGT GTGATTTGCCTTCTAGAACAGTAGACACGAAACAGGCTCAGGAGT |
| sgRNA for K104Q mouse model | 5’- TGAAGATGTGCCTATGGTCC-3’ |
